# The unfolded protein response transcription factor XBP1s ameliorates Alzheimer’s disease by improving synaptic function and proteostasis

**DOI:** 10.1101/2022.06.21.496869

**Authors:** Claudia Duran-Aniotz, Catalina Rivera-Krstulovic, Natalia Poblete, Álvaro O. Ardiles, Mei Li Díaz, Carleen Mae P. Sabusap, Yannis Gerakis, Felipe Cabral Miranda, Javier Diaz, Matias Fuentealba, Ernesto Muñoz, Sandra Espinoza, Gabriela Martinez, Gabriel Quiroz, Giovanni Tamburini, Danilo B. Medinas, Darwin Contreras, Ricardo Piña, Mychael V. Lourenco, Felipe C. Ribeiro, Sergio T. Ferreira, Carlos Rozas, Bernardo Morales, Lars Plate, Christian Gonzalez-Billault, Adrian G. Palacios, Claudio Hetz

## Abstract

Alteration in the buffering capacity of the proteostasis network is an emerging feature of Alzheimer’s disease (AD), highlighting the occurrence of endoplasmic reticulum (ER) stress. The unfolded protein response (UPR) is the main adaptive pathway to cope with protein folding stress at the ER. Inositol requiring enzyme-1 (IRE1) is an ER-located kinase and endoribonuclease that operates as a central ER stress sensor, enabling the establishment of adaptive and repair programs through the control of the expression of the transcription factor X-Box binding protein 1 (XBP1). A polymorphism in the XBP1 promoter has been suggested as a risk factor for AD. To artificially enforce the adaptive capacity of the UPR in the AD brain, we developed strategies to express the active form of XBP1 in neurons using preclinical models. Overexpression of an active form of XBP1 in the nervous system using transgenic mice significantly reduced the load of amyloid deposits in the cerebral cortex and hippocampus and preserved synaptic and cognitive function. Moreover, local delivery of XBP1 into the hippocampus of an AD mice using adeno-associated vectors improved long-term potentiation, memory performance, and dendritic spine density. Quantitative proteomics of the hippocampus revealed that XBP1 expression corrects a large proportion of the alterations observed in the 5xFAD model, restoring the levels of several synaptic proteins and factors involved in actin cytoskeleton regulation and axonal growth. Our results illustrate the therapeutic potential of targeting UPR-dependent gene expression programs as a strategy to ameliorate AD features and sustain synaptic function.

## Introduction

Alzheimer’s disease (AD) is the most common form of dementia in the elderly, described by progressive synaptic dysfunction, leading to neurodegeneration and cognitive impairment [1, 2]. AD is characterized by the abnormal deposition of protein aggregates in the brain formed by amyloid β and hyper-phosphorylated Tau [3]. Although aging is the main risk factor for developing AD, its functional relationship with the emergence of AD features remains poorly defined. Proteostasis (homeostasis of proteins) is maintained by the dynamic integration of pathways that mediate the synthesis, folding, degradation, quality control, trafficking, and targeting of proteins [4]. The decay in the buffering capacity of the proteostasis network has been pointed out as a primary hallmark of aging [5, 6], a phenomenon that may contribute to AD pathogenesis. One of the central nodes of the proteostasis network altered in AD is the endoplasmic reticulum (ER) [7, 8], a main subcellular compartment involved in protein folding and quality control. In addition, ER stress has been widely associated with the occurrence of a variety of age-related neurodegenerative diseases, including Parkinson’s disease, frontotemporal dementia, amyotrophic lateral sclerosis (ALS), and Huntington’s disease, among other pathological conditions [8–10].

To cope with ER stress, cells activate an evolutionarily conserved pathway known as the unfolded protein response (UPR), which aims to re-establish proteostasis. The UPR reinforces many aspects of the secretory pathway, including protein folding and synthesis, trafficking, degradation, and quality control mechanisms [11, 12], whereas chronic ER stress results in neurodegeneration and cell death [13, 14]. The most conserved UPR signaling branch is initiated by the ER stress transducer Inositol-requiring enzyme 1α (here referred to as IRE1). IRE1 catalyzes the unconventional splicing of the mRNA encoding XBox binding protein-1 (XBP1), excising a 26-nucleotide intron [15–17]. This processing event shifts the XBP1 mRNA reading frame, resulting in the expression of an active transcription factor, termed XBP1s, that operates as a master regulator of UPR transcriptional responses [18, 19]. IRE1 also signals as a scaffold by interacting with different signaling proteins [20] and through degradation of a cluster of mRNAs by a process termed regulated IRE1-dependent decay (RIDD) [21–23]. RIDD is proposed to control several target genes involved in inflammation, apoptosis, DNA repair, and other biological processes [24]. Thus, IRE1 is a central regulator of adaptive and pro-apoptotic programs, thus controlling cell fate under ER stress.

The presence of ER stress markers in the brain has been correlated with the progression of AD neuropathology [25]. Furthermore, a genome-wide association study identified a polymorphism in the XBP1 promoter as a risk factor for AD in the Chinese population [26]. Functional studies in fly models of AD revealed a neuroprotective activity of XBP1s against amyloid β and Tau [27, 28]. It was also reported that the behavioral impairment triggered by amyloid β in *C. elegans* is prevented by overexpression of XBP1s, whereas knocking down *Xbp1* in worms exacerbated amyloid β pathogenesis [29, 30]. Other studies reported that XBP1s expression increases amyloid β clearance [29–31], thus suggesting multiple mechanisms of neuroprotection [29]. Similarly, XBP1s overexpression in worms attenuates Tau toxicity, possibly involving Tau degradation [32]. Interestingly, the benefits of ectopically expressing XBP1s on experimental AD pathogenesis were proposed to be attenuated as animals age in *D. melanogaster* models, consistent with the idea that the activity of the UPR declines with age [29]. In contrast, other studies suggested that ablation of *Xbp1* expression can delay the paralysis triggered by amyloid β *in C. elegans* associated with the upregulation of protective macroautophagy [31]. In agreement with this concept, ablation of *Xbp1* expression in the brain protects against ALS and Huntington’s disease, involving compensatory changes that shift the proteostasis network toward increased macroautophagy levels [33, 34].

Although IRE1 and XBP1s are typically considered a linear pathway, we recently uncovered an unexpected role of IRE1 in AD pathogenesis using mouse models [25]. Despite initial expectations that IRE1 signaling might protect against AD, genetic ablation of the RNase domain of IRE1 in the central nervous system led to a significant decrease in amyloid β deposition in the brain of AD mice, and restored cognitive function and synaptic plasticity [25]. The protective effects of IRE1 deficiency correlated with increased clearance of amyloid precursor protein (APP) [25]. However, local expression of XBP1s in the hippocampus using lentiviral-mediated delivery was shown to rescue cognitive function in AD models associated with the upregulation of genes involved in synaptic plasticity [35]. Consistent with these findings, we previously reported that XBP1s has a physiological function in the nervous system at basal levels to improve synaptic function and memory performance [36]. Interestingly, an unbiased screening to identify the regulatory network governed by XBP1s in non-neuronal cells uncovered a cluster of AD-related genes involved in amyloid β biogenesis [19], and cell culture studies linked the activity of XBP1 to APP metabolism [37, 38]. The activation of XBP1 has been shown to occur transiently in AD model mice [37], and shows a marked downregulation in the brain of AD cases compared to controls [37], suggesting a loss of neuroprotective activity aging progresses.

Here we developed strategies to improve ER proteostasis in AD by expressing the active form of XBP1 in the mammalian brain. We show that XBP1s expression significantly diminishes the cognitive decline observed in experimental AD using neuronal-specific transgenic mice and gene therapy with recombinant viruses. Furthermore, XBP1s expression has a substantial effect in reducing the load of amyloid β in AD mice. To gain mechanistic insight into the positive effects of XBP1 in AD, we performed a quantitative proteomic analysis of hippocampal tissue followed by functional enrichment analysis. We identified a cluster of proteins implicated in actin dynamics and axonal growth altered in the AD model and fully corrected by XBP1s expression. Our results suggest that therapeutic strategies to enhance the activity of the adaptive UPR branches may improve synaptic function, reduce abnormal protein aggregation and delay cognitive decline in AD.

## Materials and methods

### Generation of a transgenic mouse model of AD overexpressing XBP1s in the nervous system

XBP1s overexpressing mice (Tg^XBP1s^) were described before [36]. These animals were crossed with the AD model 5xFAD animals (formerly JAX Stock No. 008730). 5xFAD mice overexpress two transgenes, which have three mutations in the human APP gene (Swedish, Florida, and London Familial Alzheimer’s Disease), and two in the human Presenilin-1 (PSEN1) gene (M146L+L286V). These animals begin to accumulate amyloid β aggregates at 2 months. Increased senile plaques, synaptic degeneration, gliosis, and cognitive impairment, are observed between 4 and 5 months of age [39]. Animals were housed in groups of a maximum of five in individually ventilated cages under standard conditions (22 °C, 12 h light-dark cycle), receiving food and water ad libitum. All animal manipulations were carried out according to standard regulations and approved by the Animal Welfare Committee of the Faculty of Medicine of the Universidad de Chile, Chile. Mice were euthanized by CO_2_ inhalation, and all efforts were made to minimize suffering. The left hemispheres were frozen at −80°C for biochemical analyses, whereas the right hemispheres were stored for histological studies.

### Behavioral studies

Cognitive impairment was measured using different behavioral tests, including the Barnes Maze (BM), Morris Water Maze (MWM), and Y Maze (YM). Briefly, the BM test consists of a circular surface containing 40 holes. Among them, only one is equipped with a hiding chamber. Mice were placed on the maze surface and were allowed to explore it for 1 minute. Mice were stimulated with sound to stress them in looking for the hiding chamber. Room environments had spatial cues for mouse orientation. This training procedure was performed four times a day for four consecutive days per animal. On the fifth day, memory was evaluated by measuring the time each animal takes to find the chamber, the time spent in the target quadrant, and the number of errors made by each animal before arriving at the objective. Errors and time to the objective were measured when the animal’s nose was entirely inside one hole. On the 5th day, short-term memory (STM) was measured. On day 12, long-term memory (LTM) was assessed. The MWM was performed as previously described [25]. In this assessment, animals learn to swim to a hidden platform under the water. Mice at 6 or 9-10 months of age were placed in the pool and allowed to explore it for 1 min. The testing room was equipped with spatial cues for orientation. This training procedure was performed six times a day for four consecutive days per animal. On day five, the platform was removed from the pool to measure the animals’ time spent in the target quadrant until the animal found the platform. Additionally, each animal’s time spent finding the platform was also measured during the training phase. Learning performance was measured as total latency on the fifth day. The Spatial Working Memory was analyzed using a Y-maze protocol. The Y-maze array consists of three arms (labeled as A, B, and C) that radiates from the center in the shape of Y. The behavior task was initiated by placing the mouse in the center of the Y, which allows free access to three arms. The animal behavior was recorded for 8 minutes, and the number of entries in each arm and the sequence of entrances were measured. A “Spontaneous Alternation” is defined as a set of three consecutive arm choices without a repeated entry (for example, ABC, BCA, CBA). A spontaneous alternation score was calculated using the formula: #alternances ÷ (number of total entries - 2) × 100). All behavior analysis were performed employing the commercial software for video tracking Any-Maze.

### Electrophysiological analysis

Hippocampal slices were prepared as previously reported [25, 36]. 6 to 9-month-old mice were deeply anesthetized with isoflurane, and their brains were quickly removed. Slices (350 μm) were dissected in ice-cold dissection buffer using a vibratome (Vibratome 1000 plus, Ted Pella Inc., CA, USA). Synaptic responses were evoked by stimulating the Schaffer collaterals with 0.2 ms pulses delivered through concentric bipolar stimulating electrodes and recorded extracellularly in the stratum radiatum of the CA1 subfield as described [25, 36]. Long-term potentiation (LTP) was induced by four-theta burst stimulation (10 trains of four pulses at 100 Hz; 5 Hz inter-burst interval) delivered at 0.1 Hz. LTP magnitude was calculated as the average (normalized to baseline) of the responses recorded 50-60 min after conditioning stimulation.

### Histological analysis

Fixed brains were collected in serial coronal sections either on a freezing cryostat or embedded in paraffin and processed on a microtome at 25-μm-thick or 12-μm-thick 12 serial slices (10 sections/stain/animal) from lambda 0 to lambda −4 mm, respectively. For the paraffin sections, a dehydration process was done after the staining. After formic acid-induced epitope retrieval, primary antibody 4G8 was incubated overnight at a 1:1000 dilution at room temperature (RT) (Biolegend, San Diego, CA). HRP-linked secondary goat anti-mouse antibody at a 1:1000 dilution (Invitrogen) was incubated for 2 h at RT. Peroxidase reaction was visualized using DAB Kit (Vector) following the manufacturer’s instructions. Finally, sections were dehydrated in ascendant ethanol, cleared in xylene, and coverslipped with DPX mounting medium (Innogenex, San Ramon, CA). For fibrillar Aβ quantification, sections were incubated in Thioflavin-S (ThS) solution (0.025% in 50% ethanol) for 10 min after defrosting. Sections were coverslipped with DPX mounting medium (Innogenex, San Ramon, CA). Astrogliosis was visualized after staining with rabbit Monoclonal Anti-Glial Fibrillary Acidic Protein (GFAP) antibody at a 1:1000 dilution (Abcam, Cambridge, MA). All samples were analyzed on an inverted epifluorescent microscope (Olympus IX71), and quantification analysis was performed using the ImageJ software.

### ELISA quantification of amyloid β species

The dissected brain was separated into cortical and hippocampal areas to generate the brain homogenates (BH). To measure Aβ_42_ levels, the BH was processed using a previously described serial extraction protocol [25, 40, 41]. 10% BH (cortex and hippocampus) were centrifuged in L100K ultracentrifuge tubes (Beckman-Coulter, Brea, CA) at 32600 r.p.m for 1 hr at 4°C in a 42.2 Ti rotor. Supernatants were collected, and pellets were resuspended in 70% Formic Acid (Fisher Scientific, Waltham, MA). Then, samples were centrifuged for 30 min, and supernatants were collected. To neutralize the samples, formic acid fractions were diluted on 1 M Tris Buffer pH 11 (Sigma-Aldrich, St. Louis, MO) 20 folds to neutralize the samples. ELISA was used to measure the levels of Aβ_42_ in the brain (kit KHB3442, Invitrogen, Carlsbad, CA) following the manufacturer’s instructions. Finally, samples were measured using an ELISA reader (EL800 BIO-TEK, BioTek, Winooski, VT) at 450 nm.

### Amyloid β oligomer preparations

Amyloid β oligomers (Aβo) were prepared weekly from synthetic Aβ_1–42_ (California Peptide, Salt Lake City, UT, USA), and were routinely characterized by size-exclusion chromatography (SEC-HPLC) under nondenaturing conditions and, occasionally, by Western immunoblots and transmission electron microscopy, as previously described [42, 43]. Briefly, the peptide was dissolved in HFIP to 1 mM and stored as a dried film at −80 °C after solvent evaporation. The film was resuspended in DMSO to a final concentration of 5 mM and thoroughly vortexed. The solution was then diluted in ice-cold PBS to 100 mM and left at 4°C overnight. The solution was centrifuged at 14,000 g for 10 min at 4°C to remove insoluble aggregates (protofibrils and fibrils), and the supernatant containing Aβo was collected. Protein concentration was determined using the BCA assay (Thermo-Pierce). Intracerebroventricular (i.c.v.) infusion of Aβo in mice was performed as described [44–46].

### Stereotaxic injections

The adeno-associated vector used was produced as described [47]. Forty-five-day-old 5xFAD mice were anesthetized using isoflurane and fixed to a mouse stereotaxic frame (David Kopf Instruments). Bilateral injections of 2.5 μL of AAV-XBP1s or AAV-Mock virus were performed at a single point in the hippocampal region using a 5 μl Hamilton syringe (Hamilton) using the following coordinates: AP: −1.8 mm, ML: 1.8 mm y DV: −1.8 mm. The injection was conducted at a rate of 0.5 μl/min, and the needle was left in place for 5 min before retraction.

### Dendritic spine imaging

Brain slices were cut at 25 mm thickness in a cryostat. AAV-GFP fluorescence was previously confirmed in injected animals to validate viral transduction in an inverted fluorescence microscope and then imaged in a confocal microscope Nikon Eclipse T1 at 60x magnification with an additional digital zoom of 3x. Similar regions were compared within each animal (CA1 region, spines in primary and secondary dendrites between the stratum radiata and the pyramidal layer, AP: 1.9-2.1 from the bregma). Z-stacks were acquired in 0.5 mm slices, laser intensity at 0.5-1%, and 12.5us/pixel at 1024×1024 resolution. Z-stacks were then summed using ImageJ software with total maximum intensity to generate one stacked 8-bit image. The number of spines was manually quantified in scaled images and divided by the dendrite length analyzed.

### Quantitative Proteomic Analysis

Hippocampal tissue of WT, Tg^XBP1s^, 5xFAD, and Tg^XBP1s^/5xFAD mice were homogenized in TEN buffers as described above. 20 μg of lysate was precipitated with chloroform/methanol for each sample. Samples for mass spectrometry analysis were prepared as described [48]. Air-dried pellets were resuspended in 1% RapiGest SF (Waters) and diluted to final volume in 100 mM HEPES (pH 8.0). Proteins were reduced with 5 mM Tris (2-carboxyethyl) phosphine hydrochloride (Thermo Fisher) for 30 min and alkylated with 10 mM iodoacetamide (Sigma Aldrich) for 30 min at room temperature in the dark. Proteins were digested for 18 h at 37°C with 0.5 μg trypsin (Promega). After digestion, the peptides from each sample were reacted for 1 h with the appropriate tandem mass tag (TMT) isobaric reagent (Thermo Fisher) in 40% (v/v) anhydrous acetonitrile and quenched with 0.4% ammonium bicarbonate for 1 h. Samples with different TMT labels were pooled and acidified with 5% formic acid. Acetonitrile was evaporated on a SpeedVac, and debris was removed by centrifugation for 30 min at 18,000g. MudPIT microcolumns were prepared as described [49]. LC-MS/MS analysis was performed using a Q-Exactive HF mass spectrometer equipped with an Ultimate 3000 nLC 1000 (Thermo Fisher). MudPIT experiments were performed by 10 μl sequential injections of 0, 10, 20, 30, …, 100% buffer C (500 mM ammonium acetate in buffer A) and a final step of 90% buffer C/10% buffer B (100% acetonitrile, 0.1% formic acid, v/v/v) and each step followed by a gradient from buffer A (95% water, 5% acetonitrile, 0.1% formic acid) to buffer B. Electrospray was performed directly from the analytical column by applying a voltage of 2 kV with an inlet capillary temperature of 275°C. Data-dependent acquisition of MS/MS spectra was performed with the following settings: eluted peptides were scanned from 300 to 1800 m/z with a resolution of 120,000. The top 15 peaks for each full scan were fragmented by HCD using a normalized collision energy of 30%, isolation window of 2.0 m/z, a resolution of 30,000, ACG target 1e5, maximum IT 60 ms, and scanned from 100 to 1800 m/z. Dynamic exclusion was set to 10 s. Proteome Discoverer 2.2 (ThermoFisher) performed peptide identification and protein quantification. Spectra were searched using SEQUEST against a UniProt mouse proteome database. The database was curated to remove redundant protein and splice-isoforms, and common contaminants were added. Searches were carried out using a decoy database of reversed peptide sequences using Percolator node for filtering and the following settings: 50 ppm peptide precursor tolerance, 6 amino acid minimum peptide length, trypsin cleavage (unlimited missed cleavage events), static Cys modification of 57.0215 (carbamidomethylation), and static N-terminal and Lys modification of 229.1629 (TMT sixplex), FDR 0.01, 2 peptide IDs per protein. TMT reporter ion intensities were normalized based on total peptide abundance in each channel. Subsequently, a common pooled sample calculated TMT ratios for each identified protein. Finally, the reference-normalized TMT intensities were compared between WT (n = 5), Tg^XBP1s^ (n = 5), 5xFAD (n = 5) and Tg^XBP1s^/5xFAD (n = 5). Statistical significance was assessed by a twotailed unpaired t-test using the FDR approach [50] and Q = 1% in Graphpad Prism. Enrichment for gene ontology (GO) terms was tested in EnrichR [51, 52].

To identify changes in the proteome we fitted a linear model of proteomic levels versus genotype and the samples run for each protein and calculated the summary statistics using a moderated t-test implemented in the R package limma [53]. Correlations between results of the comparative proteomics analysis were performed using Spearman’s method on significance scores calculated as the sign of the fold changes (1 for positive values and - 1 for negative values) multiplied by the logarithm of the p-values. The gene set enrichment analysis was performed using proteins ranked by significance scores as input for the function gseGO from the R package ClusterProfiler [54]. P-values for the enriched GO terms were adjusted for multiple testing using the Benjamini-Hochberg method [55]. The statistical significance of the overlaps was calculated using a Fisher’s Exact Test [56]. To determine the genes leading the top enriched processes, we performed a leading-edge analysis using the R package fgsea [57]. To search proteomics datasets of the 5xFAD mouse model, AD, and human aging, we used the PRIDE database (see raw values in Supplementary Table 1) [58]. We normalized the log2 fold changes using quantile normalization to compare the effect sizes across datasets.

### Statistical Analyses

Data are expressed as means ± SEM. After confirming normal distribution with the Skewness/Kurtosis statistic test, Student’s t-test was used to analyze differences in histological and biochemical analysis of Aβ and APP. For behavioral studies, one-way ANOVA followed by Tukey’s Multiple post-test or Newman-Keuls Multiple post-test measured significant differences. Two-way ANOVA followed by Bonferroni post-test was used in electrophysiological and behavioral experiments. Statistical analyses were performed using Graph Pad Prism 5.0 software. Statistical differences were considered significant for values of *p* < 0.05.

## Results

### Overexpression of XBP1s in the brain reduces amyloid β load in AD mice

To determine the consequences of enforcing adaptive UPR responses in the AD brain, we first used a transgenic mouse model generated by our laboratory that overexpresses XBP1s through the control of the Prion promoter (Tg^XBP1s^) [36]. These animals were crossed with the 5xFAD model, which expresses a combination of five human mutations in APP and PSEN1 genes [39] (Supplementary Fig. S1).

We next determined the effects of overexpressing XBP1s in the progression of key AD features. We evaluated amyloid β deposition in the brain of Tg^XBP1s^ crossed with 5xFAD animals (termed Tg^XBP1s^/5xFAD) of experimental mice at 6 and 8 months of age (Fig. 1). Immunohistochemical analysis of amyloid β content using the 4G8 antibody showed a significant reduction in the cerebral cortex of Tg^XBP1s^/5xFAD mice compared to 5xFAD littermate control animals (Fig. 1a). Similar results were obtained when hippocampal tissue was analyzed (Fig. 1d). Overall, Tg^XBP1s^/5xFAD mice showed a 30-40% reduction in amyloid β load in both brain regions compared with 5xFAD at the same age (Fig. 1b and 1e). These results were also confirmed by measuring the number of amyloid β plaques per area in both cortical and hippocampal regions. We observed a decrease in the number of amyloid deposits when XBP1s was overexpressed (Fig. 1c and 1f). Then, we stained brain tissue with thioflavin S (ThS) to evaluate fibrillar amyloid deposits. Expression of XBP1s also reduced ThS-reactive deposits in the cortex and hippocampus of 5xFAD mice at 8 months old (Fig. 1g).

**Fig 1.**
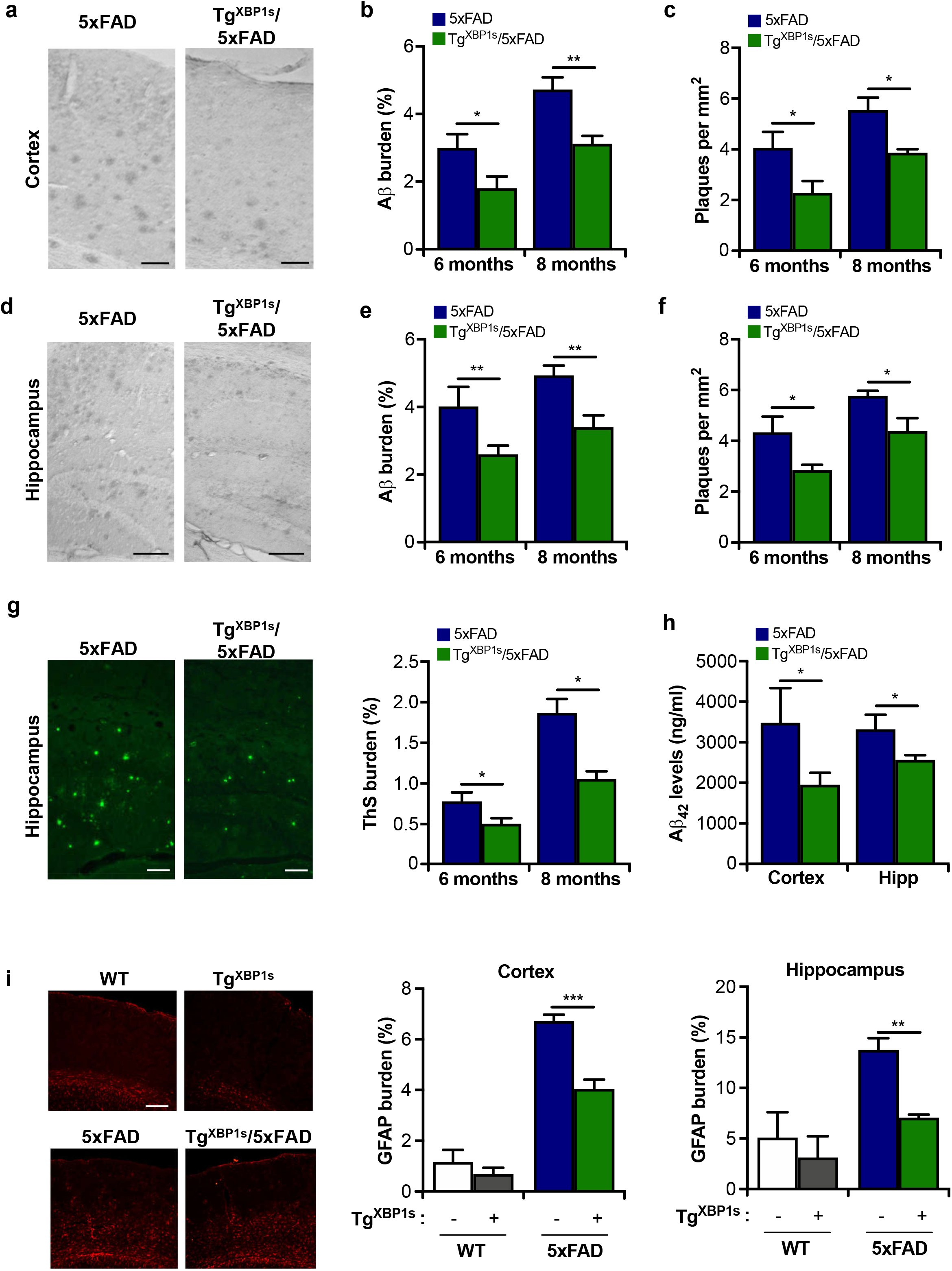
Overexpression of XBP1s in the nervous system reduces amyloid β levels. Tg^XBP1s^ were crossed with 5xFAD mice, and histopathological and biochemical analyses were performed in brain. Representative images of amyloid β (Aβ) deposits in cortical **(a)** and hippocampal **(d)** areas of 6-month-old 5xFAD and Tg^XBP1s^/5xFAD animals. Scale bar: 100 μm. The burden of amyloid deposits and the number of deposits per mm^2^ were quantified in serial brain slices (10 sections/stain/animal) of the cortex **(b,e)** and hippocampus **(c,f)** of 5xFAD and Tg^XBP1s^/5xFAD mice at 6 and 8 months of age, respectively. **(g)** Fibrillar amyloid β deposits were stained and quantified in hippocampal areas of 5xFAD and Tg^XBP1s^/5xFAD animals at 6 and 8 months of age using ThS. Scale bar: 100 μm. **(h)** Insoluble Aβ_42_ levels were quantified in hippocampal homogenates after a serial extraction protocol (see methods) followed by detection using human-specific ELISA assay. **(i)** Representative images and quantification of immunofluorescence of GFAP in cortical and hippocampal areas of 5xFAD and Tg^XBP1s^/5xFAD animals. Scale bar: 100 μm. Values are expressed as means ± SEM. 5xFAD (n = 4-10) and Tg^XBP1s^/5xFAD (n = 4-8) animals. **(g)** Quantification of immunofluorescence of GFAP in cortical and hippocampal areas of 5xFAD and Tg^XBP1s^/5xFAD animals. Data were analyzed by Student’s t-test. *: p < 0.05; **: p < 0.01; ***: p < 0.001.

Because amyloid β oligomers are proposed as a causative agent of synaptic dysfunction and behavioral impairment in AD [59], we monitored the relative levels of soluble and insoluble amyloid β species. We performed serial extractions with detergents followed by amyloid β quantification in cortical and hippocampal brain homogenates using a specific human ELISA assay to detect the amyloid β_42_ isoform [25]. Tg^XBP1s^/5xFAD animals contained lower aggregated/fibrillar (formic acid insoluble) amyloid β_42_ species in the cortex (Fig. 1h). The soluble species showed similar levels following serial extraction in 5xFAD animals (Supplementary Fig. S2).

We then characterized other typical neuropathological alterations of AD in our experimental groups by monitoring astrocyte activation in the brain. Immunofluorescence analysis of GFAP staining indicated that astrogliosis was significantly induced in the cortex and hippocampus of 5xFAD mice at 8 months (Fig. 1i). Overexpression of XBP1s significantly attenuated the content of reactive astrocytes in both brain regions (Fig. 1i). These results indicate that XBP1s overexpression significantly reduced two central neuropathological features of experimental AD, amyloid β deposition and gliosis.

### XBP1s overexpression in the brain ameliorates cognitive impairment in AD mice

We and others previously reported that XBP1s expression in the brain enhances synaptic function and improves learning and memory-related processes [35, 36]. To monitor the possible effects of overexpressing XBP1s on the behavioral performance of 5xFAD, we evaluated spatial memory acquisition using the Barnes Maze test (Fig. 2a). In line with previous studies [60, 61], 5xFAD animals spent significantly less time than non-transgenic mice in the target quadrant (Fig. 2b). Remarkably, Tg^XBP1s^/5xFAD mice increased total time spent in the target quadrant compared to 5xFAD control animals, showing significant prevention of the memory impairment (Fig. 2b).

**Fig 2.**
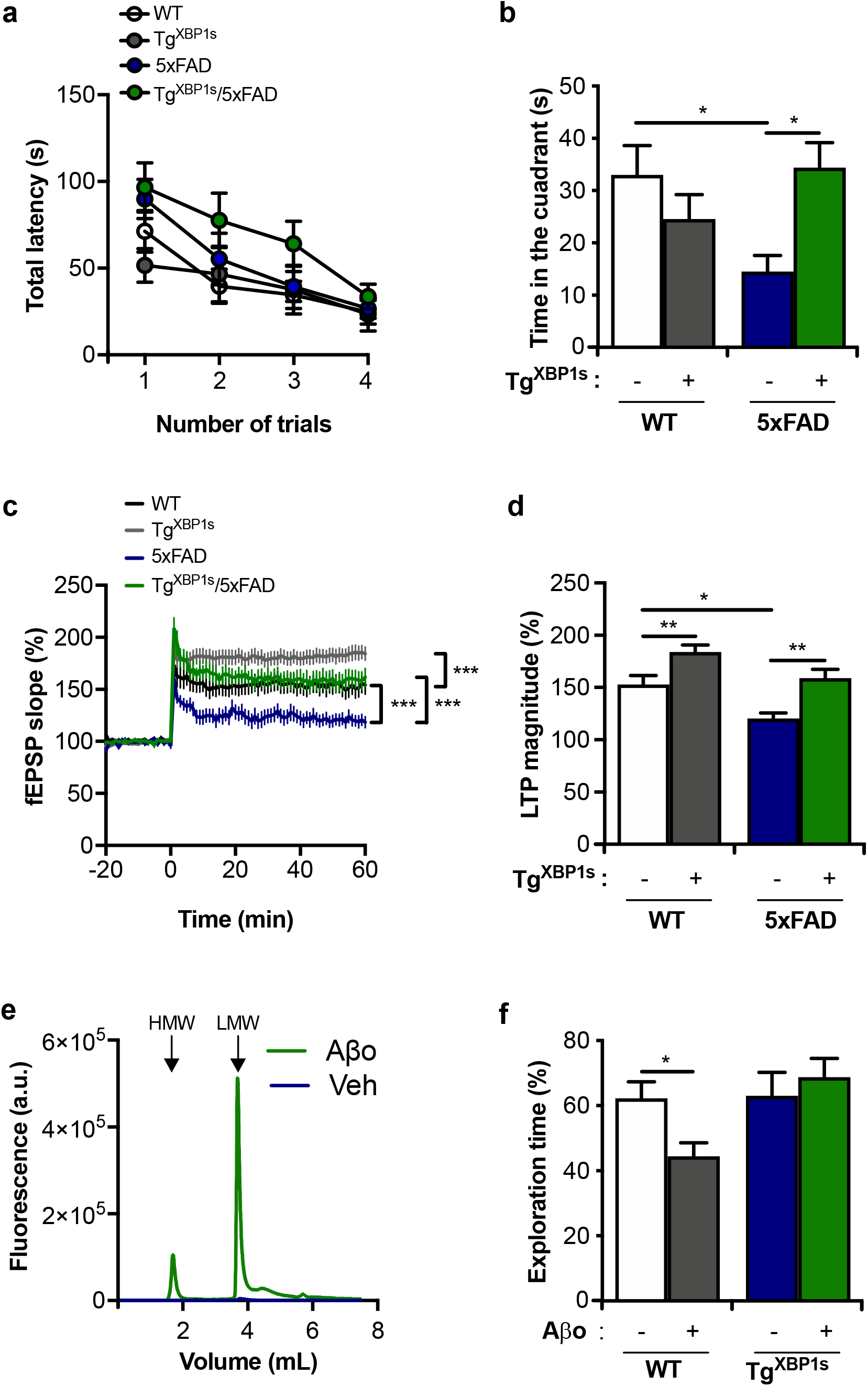
XBP1s overexpression in the nervous system attenuates cognitive deficits and synaptic alterations of 5xFAD mice. WT (n = 15), Tg^XBP1s^ (n = 8), 5xFAD (n = 14), Tg^XBP1s^/5xFAD (n = 14) mice were analyzed using the Barnes maze behavioral test at 8 months of age. Learning performance was measured as total latency during the training phase **(a)**, and the time in the correct quadrant at the test day was evaluated **(b)**. The excitatory synaptic transmission was analyzed by long-term potentiation (LTP) in hippocampal slices from animals presented in **a**. **(c)** Input-output relationship between fEPSP slope and fiber volley amplitude was monitored (n = 20-40 slices from 4-5 animals per group). **(d)** Representative traces of the field excitatory postsynaptic potentials (fEPSPs) magnitude of hippocampal LTP induced by theta-burst stimulation (TBS) protocol is shown (n = 17-36 slices from 4-6 animals per group). **(e)** *In vitro* generated amyloid β oligomers (Aβo) were analyzed by size exclusion chromatography to assess its oligomerization state and measure low molecular (LMW) and high molecular weight (HMW) species. **(f)** Percentage of exploration time of the novel object in the NOR test was measured on day 7 after i.c.v. injection of amyloid β oligomers (Aβo) in 3 month-old Tg^XBP1s^ and WT mice. Data values were expressed as mean ± SEM. Data from **a** and **c** were analyzed by two-way ANOVA followed by Tukey’s multiple comparison post-test. Data from **b**, **d**, and **e** were analyzed by one-way ANOVA followed by Tukey’s multiple comparison post-test. *:*p* < 0.05; **:*p* < 0.01; ***: *p* < 0.001.

Next, to determine whether the expression of XBP1s could prevent synaptic dysfunction in the context of experimental AD, we measured long-term potentiation (LTP) in the hippocampus, a long-lasting form of synaptic plasticity. We recorded glutamatergic transmission evoked by Schaffer’s collaterals stimulation to monitor field excitatory postsynaptic potentials (fEPSPs) in the CA1 region [62]. We confirmed drastic decrease in the slope of fEPSP in brain tissue of 5xFAD (Fig. 2c). Strikingly, overexpression of XBP1s in the nervous system prevented LTP impairment in 5xFAD mice and significantly improved it compared with WT (Fig. 2d). Thus, the beneficial effects provided by XBP1s to the cognitive function of AD mice were associated with prevention of synaptic dysfunction at the electrophysiological level.

To validate the possible protective effects of XBP1s expression on AD, we used an alternative disease model based on intracerebroventricular infusion of amyloid β oligomers (AβOs) [44, 63]. AβOs were prepared weekly from synthetic amyloid β_1–42_ and revealed a mixture of low- and high-molecular weight amyloid β species, characterized by size-exclusion chromatography (SEC-HPLC) (Fig. 2e). Seven days after the injection of AβOs, we assessed cognitive alterations using the novel object recognition test, a task previously shown to be impaired in wild-type mice injected with AβO preparations [44, 45]. We assessed cognitive impairments in wild-type and Tg^XBP1s^ animals after injection of AβOs compared to control mice injected with a saline solution (Fig. 2f). Strikingly, mice overexpressing XBP1s in the brain were fully protected against the adverse effects of AβO exposure, showing a behavioral performance similar to vehicle-injected control animals (Fig. 2f). Overall, these results indicate that XBP1s overexpression significantly restores synaptic function and spatial memory acquisition in mouse models of AD.

### Gene therapy to deliver XBP1s into the hippocampus of AD mice restores cognitive function

Although our results in animals overexpressing XBP1s are important to define its function as a possible disease modifier and study neuroprotective mechanisms, this approach does not prove a therapeutic potential because the transgene is expressed in all neurons since embryonic development. To evaluate the possible beneficial effects of XBP1s overexpression on AD pathogenesis with a strategy that can be translated into the clinic, we locally targeted the hippocampus using brain stereotaxis to deliver adenovirus-associated viruses (AAV) expressing XBP1s [36]. These viral particles also contained an EGFP cassette for detection. 5xFAD mice were injected with AAV-XBP1s or empty vector (AAV-Mock) at 6 weeks of age and then evaluated for behavioral performance 7 to 9 months later. Remarkably, local administration of AAV-XBP1s into the hippocampus of 5xFAD mice improved the performance in learning and memory tasks as assessed using the MWM (Fig. 3a). Similar results were obtained when memory was measured in the Barnes maze test (Fig. 3b). Importantly, analysis of the latency to the target hole during the test day also indicated that the performance of 5xTg mice injected with AAV-XBP1s was equivalent to non-transgenic animals (Fig. 3c). We confirmed an improvement of cognition by measuring spatial working memory in the Y maze, observing that the bilateral injection of AAV-XBP1s into the hippocampus improved the performance of 5xFAD mice (Fig. 3d). Thus, the local delivery of XBP1s into the hippocampus prevents the progression of AD-related cognitive decline.

**Fig 3.**
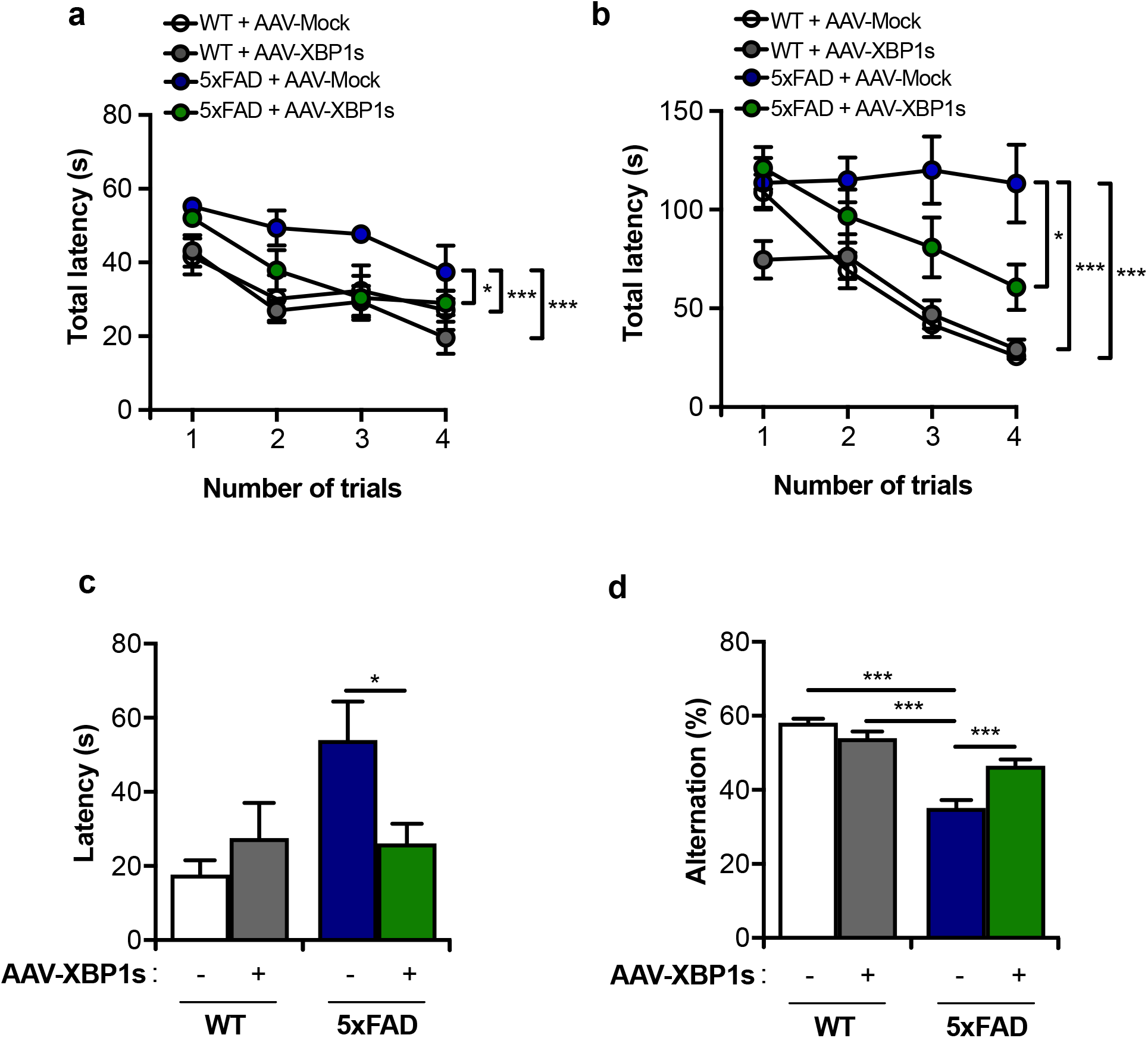
Therapeutic effects of AAV-XBP1s administration into the hippocampus of AD mice. WT and 5xFAD mice were injected with either AAV-XBP1s or AAV-Mock at 1.5 months of age and sacrificed after 8-9 months. Learning curve using the Morris Water Maze **(a)** and Barnes Maze (BM) **(b)** behavioral tests were measured. Long-term memory was measured at 12 days post-training of BM paradigm **(c)**. Percentage of Alternation of behavioral paradigm Y Maze **(d)**. Data values were expressed as mean ± SEM. WT^AVV-Mock^ (n = 7-10), WT^AVV-XBP1s^ (n = 7-10), 5xFAD^AVV-Mock^ (n = 7-10), 5xFAD^AVV-XBP1s^ (n = 7-10). Data from **c** and **d** were analyzed by one-way ANOVA followed by Tukey’s multiple comparison post-test. Data from **a** and **b** were analyzed by two-way ANOVA followed by Tukey’s multiple comparison post-test. *: *p* < 0.05; **:*p* < 0.01; ***:*p* < 0.001.

### AAV-XBP1s administration into the hippocampus of AD mice restores synaptic plasticity

As most cognitive paradigms employed here are dependent on the normal function of the hippocampus, we evaluated the distribution of dendritic spines in CA1 pyramidal cells because their density correlates with the learning capacity [64]. We injected animals with either AAV-XBP1/EFGP or AAV-Mock/EGFP to assess the morphology and content of dendritic spines utilizing EFGP fluorescence and confocal microscopy analysis., 5xFAD mice showed reduced dendritic spines per length (Fig. 4a). Consistent with our behavioral data, delivery of AAV-XBP1s into the hippocampus restored the content of dendritic spines to numbers almost comparable to littermate wild-type animals (Fig. 4a). Next, we performed electrophysiological studies in isolated hippocampal tissue from 5xFAD mice treated with AAV-XBP1s. Administration of AAV-XBP1 into the hippocampus of 5xFAD mice significantly improved fEPSP slopes (Fig. 4b). Virtually identical results were obtained when the magnitude of LTP was quantified (Fig. 4c), suggesting that the local expression of XBP1s in the hippocampus strongly improves synaptic plasticity in 5xFAD animals.

**Fig 4.**
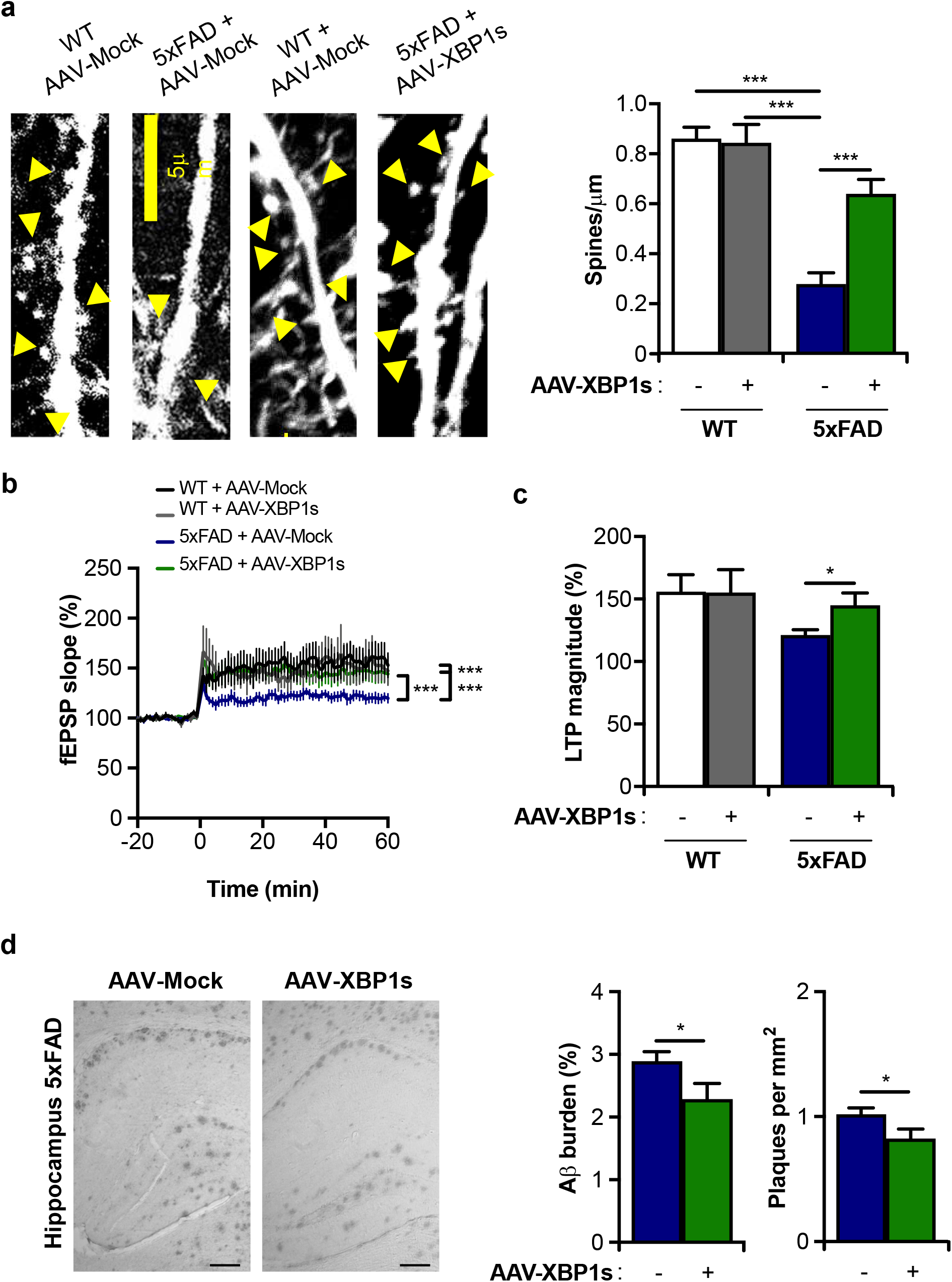
AAV-XBP1s administration into the hippocampus of 5xFAD mice improves synaptic function. **(a)** Dendritic spine density of pyramidal neurons in the CA1 region was measured in indicated experimental groups. Left panel: representative images of dendritic spines, and the quantification (right panel). Scale bar: 100 μm. **(b)** Time course of the TBS-induced LTP in slices prepared from all experimental groups and **(c)** quantification of LTP using hippocampal slices. **(d)** Representative images of amyloid deposits in the hippocampus (left panel), and quantification of amyloid β load, and the number of plaques of injected animals (right panel). Scale bar: 100 μm. Data values were expressed as mean ± SEM. WT^AVV-Mock^ (n = 7-10), WT^AVV-XBP1s^ (n = 7-10), 5xFAD^AVV-Mock^ (n = 7-10), 5xFAD^AVV-XBP1s^ (n = 7-10). Data was analyzed by one-way ANOVA followed by Tukey’s multiple comparison post-test. Data from **b** was analyzed by two-way ANOVA followed by Tukey’s multiple comparison post-test. *: *p* < 0.05; **:*p* < 0.01; ***:*p* < 0.001.

To further evaluate the consequences of ectopically expressing XBP1s on AD mice, we measured amyloid deposition after 9 months of injection of the AAV particles. Although the local injection of AAVs only transduced a portion of the hippocampus, we were able to detect a slight but significant reduction in amyloid β deposition and plaque number in animals receiving the AAV-XBP1s vector (Fig. 4d), suggesting improved proteostasis.

Finally, we injected a cohort of wild-type animals to assess the possible toxic effects of overexpressing XBP1s in the hippocampus after a long-term administration of AAV particles. Remarkably, we were able to confirm the expression of the *Xbp1s* transgene 12 months after injection, with no overall neurotoxicity (Supplementary Fig. S3a and S3b). In addition, no signs of ER stress were observed in these brain samples when we measured the mRNA levels of the PERK/ATF4 target genes *Bip* and *Chop* by real-time PCR (XBP1-independent targets) in wild-type animals (Supplementary Fig. S3c).

Altogether, these results indicate that local administration of AAV-XBP1s into the hippocampus is safe and effective in restoring synaptic plasticity and cognitive function in an animal model of AD.

### XBP1s corrects the proteomic alterations observed in the brain of 5xFAD mice

Because XBP1s-target genes are functionally diverse, this UPR factor might attenuate the progression of experimental AD by multiple mechanisms. For example, XBP1s may improve ER proteostasis by regulating canonical UPR targets, by regulating synaptic function, or by modifying other biological functions previously linked to XBP1s (i.e. energy and lipid metabolism, cell differentiation, inflammation, etc. [12, 14, 65]). To determine the major gene expression alterations in the hippocampus of AD mice and how XBP1s modify them, we performed quantitative proteomics of dissected hippocampus to compare tissues derived from Tg^XBP1s^, 5xFAD, Tg^XBP1s^/5xFAD, and littermate control animals (WT) (Fig. 5a). Interestingly, overexpression of XBP1s induced different proteomic changes in WT mice and the AD model (r = −0.02, p = 0.34), indicating that the disease context influences the impact of XBP1s overexpression on gene expression. Notably, 76% of the alterations observed in the 5xFAD model were corrected when XBP1s was overexpressed in the disease model (r = −0.71, p = 1.61e-217), suggesting global benefits at the proteomic level (Fig. 5a).

**Fig 5.**
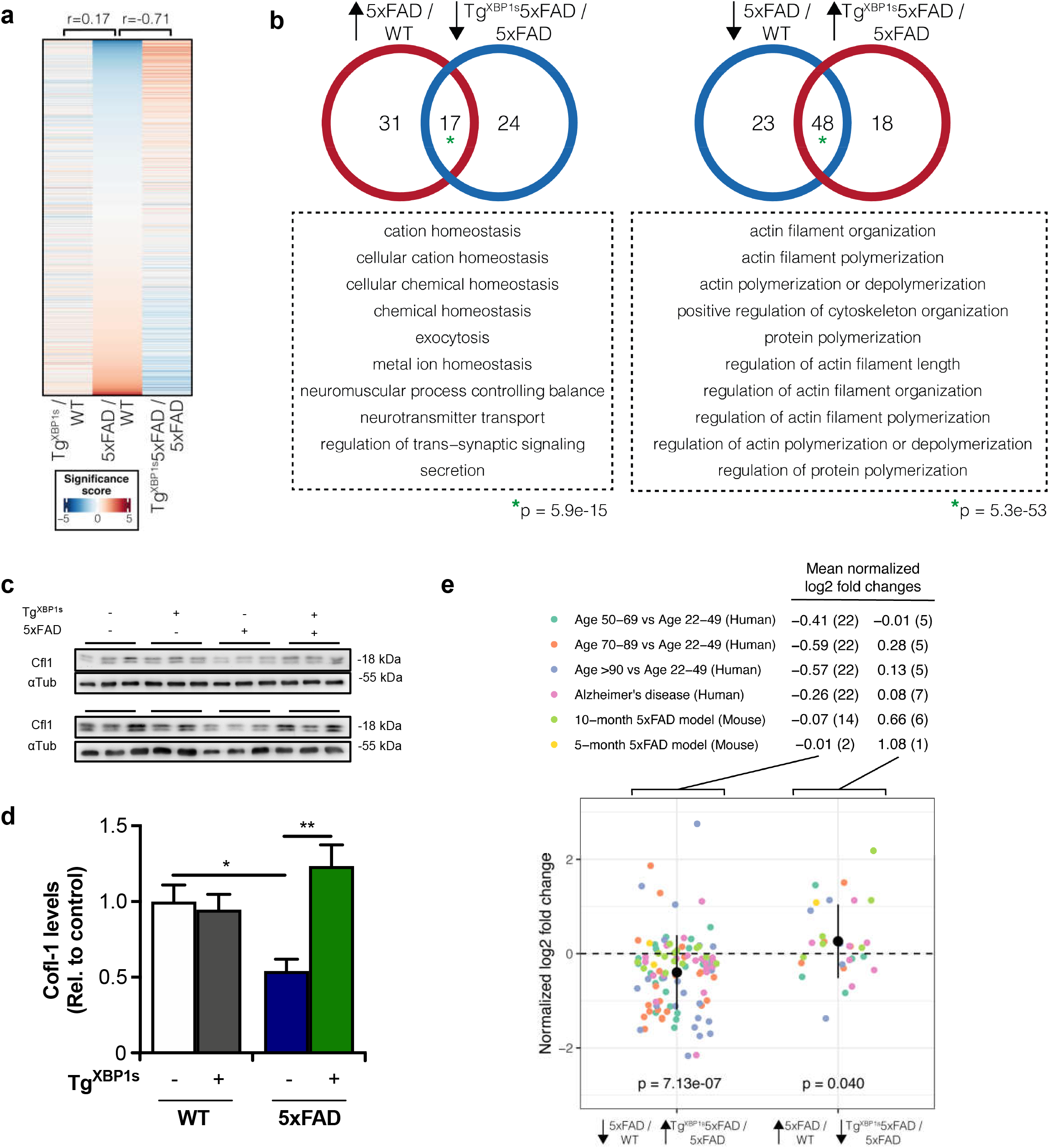
Comparative proteomics analysis of hippocampal tissue of AD mice overexpressing XBP1s. **(a)**. The significance score in the heatmap represents a combination of the statistical significance and the direction of change (see Methods). The values at the top represent Spearman’s correlation coefficient between the groups. **(b)** Overlap of the biological processes regulated in opposite directions in WT vs. 5xFAD and 5xFAD vs. Tg^XBP1s^/5xFAD mice. Top-10 enriched overlapping processes are displayed in the dashed boxes. P-values at the bottom represent the statistical significance of the overlap. **(c)** Cfl1 Western Blot and the respective quantification of hippocampal lysate from WT, Tg^XBP1s^, 5xFAD, and Tg^XBP1s^/5xFAD mice, using αTubulin as a housekeeping control. **(d)** Proteomic changes in the hippocampus of Alzheimer’s disease patients and humans at different ages for the top proteins restored by XBP1s overexpression. At the top, we show the mean fold changes for the candidate proteins in each comparison group. In parenthesis, we indicate the number of proteins measured in each dataset. The dot plot below displays the quantile normalized fold change values of the proteins in each dataset (dot color). P-values at the bottom indicate the statistical significance from a t-test with population mean equal zero.

To identify processes targeted by XBP1 in the AD model, we performed gene set enrichment analysis on proteins altered of 5xFAD mice and corrected by XBP1s overexpression. Thus, we searched for biological processes significantly down-regulated in the disease model and up-regulated after XBP1s overexpression or vice versa (Fig. 5b). In both cases, we found a statistically significant overlap between the pathophysiological processes altered in AD mice that are modified in opposite directions by XBP1s overexpression in the disease context. Biological processes associated with ion transport and cell homeostasis showed up-regulation in the AD model and were reversed upon XBP1s overexpression. At the same time, components involved in actin and the cytoskeleton regulation were downregulated in the disease model and reversed by XBP1s (Fig. 5b and supplementary Fig. S4a and S4b).

Because the major protective effects of XBP1s overexpression in 5xFAD model mice involved improvements in synaptic plasticity and memory performance, we decided to focus our proteomic validation analysis on assessing the impact on actin cytoskeleton function. The regulation of actin cytoskeleton dynamics is essential to sustain neuronal connectivity, synaptic plasticity, dendritic spine formation, and general brain function [60, 61]. We identified 23 interesting genes enriched in more than half of the top 10 biological processes (Supplementary Fig. S4a and S4b), among which we identified cofilin-1 (Cfl1) as a strong hit, a protein previously associated with neurodegenerative diseases [66]. In addition, we identified three putative UPRE elements (XBP1s DNA binding sequence) of the human *Cofilin-1* gene through the bioinformatic analysis of the proximal promoter region [59] (Supplementary Fig. S5). We confirmed the downregulation of cofilin-1 in hippocampal brain extracts derived from in 5xFAD animals using western blot analysis, a phenomenon restored in Tg^XBP1s^/5xFAD animals (Fig. 5c and 5d), validating a major hit of our proteomic analysis. We also identified 7 key proteins involved in processes linked with ion transport that increased abundance in the disease model but decreased their levels after XBP1s overexpression (Supplementary Fig. S4c and S4d)

To increase the relevance of our findings to human aging and AD, we assessed the levels of the identified genes in in publicly available proteomics datasets (Fig. 5e, supplementary Table S1). We selected 30 proteins that are altered in our AD model and corrected them by XBP1. On average, proteins with decreased abundance in the disease model and restored by XBP1s overexpression also decreased during normal human aging and in also in tissues derived from AD patients (Fig. 5e). Similarly, increased abundance was observed in AD patients and people aged 70 or higher in genes up-regulated in 5xFAD animals and down-regulated in Tg^XBP1s^/5xFAD mice. A similar consistency was observed with a previous proteomics dataset of the 5xFAD disease model [67]. To calculate the global significance of these trends, we pooled the fold change values of these proteins in all the external datasets analyzed (Fig. 5e). Overall, we observed that during human aging and AD, the proteins changed in the same direction as in our AD model (p-value for deviation from zero < 0.05), suggesting that proteins altered in the disease model and restored by XBP1s are also affected during human aging and in the brain of AD patients.

## Discussion

According to The World Alzheimer Report 2021 [68], approximately 55 million people live with dementia worldwide. AD is the most common cause of dementia and has a poorly understood etiology [1, 3]. Accumulating evidence supports the concept that impairment in the adaptive and buffer capacity of the proteostasis network is a salient feature of AD, in addition to aging, the major risk factor for developing AD [70]. We highlight the involvement of protein folding stress responses at the ER as a transversal pathological event observed in AD patient-derived tissue in addition to most animal and cellular models of the disease [7, 8, 71]. Importantly, attenuation of ER stress with pharmacological or gene therapy strategies affords strong protection in various animal models of brain diseases and thus holds promise as therapeutic approaches for human neurodegenerative diseases, including AD [9, 72–74].

XBP1 is a key transcription factor of the UPR mediating adaptive responses through the upregulation of genes in part involved in protein folding and quality control mechanisms. Here we report complementary evidence supporting a functional involvement of XBP1s in the attenuation of AD features, improving cognitive and synaptic function, and reducing the accumulation of amyloid β levels in the brain. In agreement with our results, Cissé et al. reported that brain delivery of XBP1s using lentiviruses improved synaptic plasticity and cognitive function in 3xTg-AD mice, a less aggressive AD model compared to 5xFAD, through the regulation of synaptic regulator Kalirin-7 [30]. Here we report the consequences of the bilateral hippocampal injection of AAV-XBP1s into 5xFAD at an age where the accumulation of amyloid β is already occurring. In agreement with our results, other reports suggested that the artificial enforcement of XBP1s-dependent responses using gene therapy promotes a variety of beneficial effects in other disease models, including improved dopaminergic neuron survival [75, 76], reduced mutant huntingtin aggregation [77], improved locomotor recovery after spinal cord injury [78], accelerated axonal regeneration of peripheral nerves [79], in addition to improved synaptic plasticity at basal levels [36].

The involvement of IRE1 in AD is complex possibly due to the divergent downstream signaling outputs of the pathway. We previously reported an unexpected role of IRE1 in AD pathogenesis [25] where genetic ablation of IRE1 function in the brain of 5xFAD mice significantly decreased amyloid deposition and improved synaptic function. At the mechanistic level, IRE1 deficiency reduced APP steady-state levels, and thus amyloid β production. However, in that study, we could not discriminate the differential contribution of RIDD, XBP1s, or the scaffold function of IRE1 to AD pathogenesis *in vivo*. Importantly, several RIDD targets regulate inflammation, cell migration, and metabolism, among other biological processes [23, 80, 81]. RIDD has also been associated with controlling cell fate under ER stress through caspase activation and the modulation of death receptor signaling [23, 82]. Interestingly, one of the canonical RIDD targets is collagen 6 [22], a gene protecting against AD in mouse models [83, 84]. However, the function of RIDD activity in AD remains to be determined. IRE1 also mediates the activation of JNK and NF-κB pathways [14], in addition to associating with other signaling proteins as a scaffold, a non-canonical function impacting different cellular processes, including energy metabolism [85] and cytoskeleton dynamics [86]. Thus, defining the exact contribution of IRE1 to experimental AD at the mechanistic levels deserves further investigation.

Our results suggest that XBP1s overexpression has outstanding effects in alleviating and preventing the cognitive decline observed in preclinical AD models. Local delivery of XBP1s into the hippocampus of adult animals was sufficient to restore brain function to almost normal levels. Our unbiased proteomic analysis did not reveal any clear changes in canonical proteostasis genes regulated by XBP1s in non-brain tissue. Instead, a robust cluster of genes associated with actin cytoskeleton dynamics and axonal growth were altered in the AD model, which was fully corrected by the genetic enforcement of XBP1s expression. These results uncovered a completely new function of XBP1s in the nervous system and agree with previous observations suggesting virtually no changes in the expression levels of canonical ER stress-related genes when XBP1 levels are manipulated in the brain [36]. Because our AAV-XBP1 gene therapy has partial effects on amyloid deposition but fully restored cognitive and synaptic function, we speculate that a major protective mechanism of XBP1s in AD relates to its function as a regulator of neuronal function. Since gene therapy has passed regulatory approval by the FDA and has shown outstanding efficacy in treating various diseases in clinical trials, it promises the possibility of intervene pathogenic pathways or delivering disease-modifier agents like XBP1s into the brain of AD patients. Preclinical studies are required in non-human primates to assess the safety issues related to the chronic expression of adaptive UPR mediators. In our hands, we have bred XBP1s transgenic mice for a decade without observing any evident side effects, neurotoxicity, or signs of cancer (unpublished observations). Altogether, our findings suggest that the local enforcement of XBP1s expression in the hippocampus might serve as a strategy to preserve brain function in AD and other dementias and even recover the functionality of the existing neurons undergoing synaptic impairment.

## Acknowledgments

We thank Francisco Aburto, Maria Jose Altamirano, and Javiera Ponce for technical support in animal care supervision. We thank Claudia Sepulveda and Susana Manriquez for lab managing and administrative accounting. This work was directly funded by Alzheimer’s Association Research Grant 2018-AARG-591107 (CDA), U.S. Air Force Office of Scientific Research FA9550-21-1-0096 (CH), FONDAP program 15150012, Millennium Institute P09-015-F, and Department of Defense grant W81XWH2110960 (CH). We also thank the support from ANID/FONDEF ID16I10223 and ANID/FONDECYT 1220573 (CH), 1200880 (AGP), 1161524 (BM), 3190897 (DC), 1191538 (DBM). We also thank the support ICM-P09-022-F (AOA and AGP), a binational collaborative grant from CONICYT(Chile)-CNPq(Brazil) 441921/2016-7 (CH, STF), Alzheimer’s Association AARG-D-615714 (MVL) and Serrapilheira Institute R-2012-37967 (MVL), ANID/FONDECYT 11140430 and DICYT 021843RS (CR), DICYT-Usach 021843MM, 021943MM.

## Supplementary figure legends

**Supplementary Fig S1.**
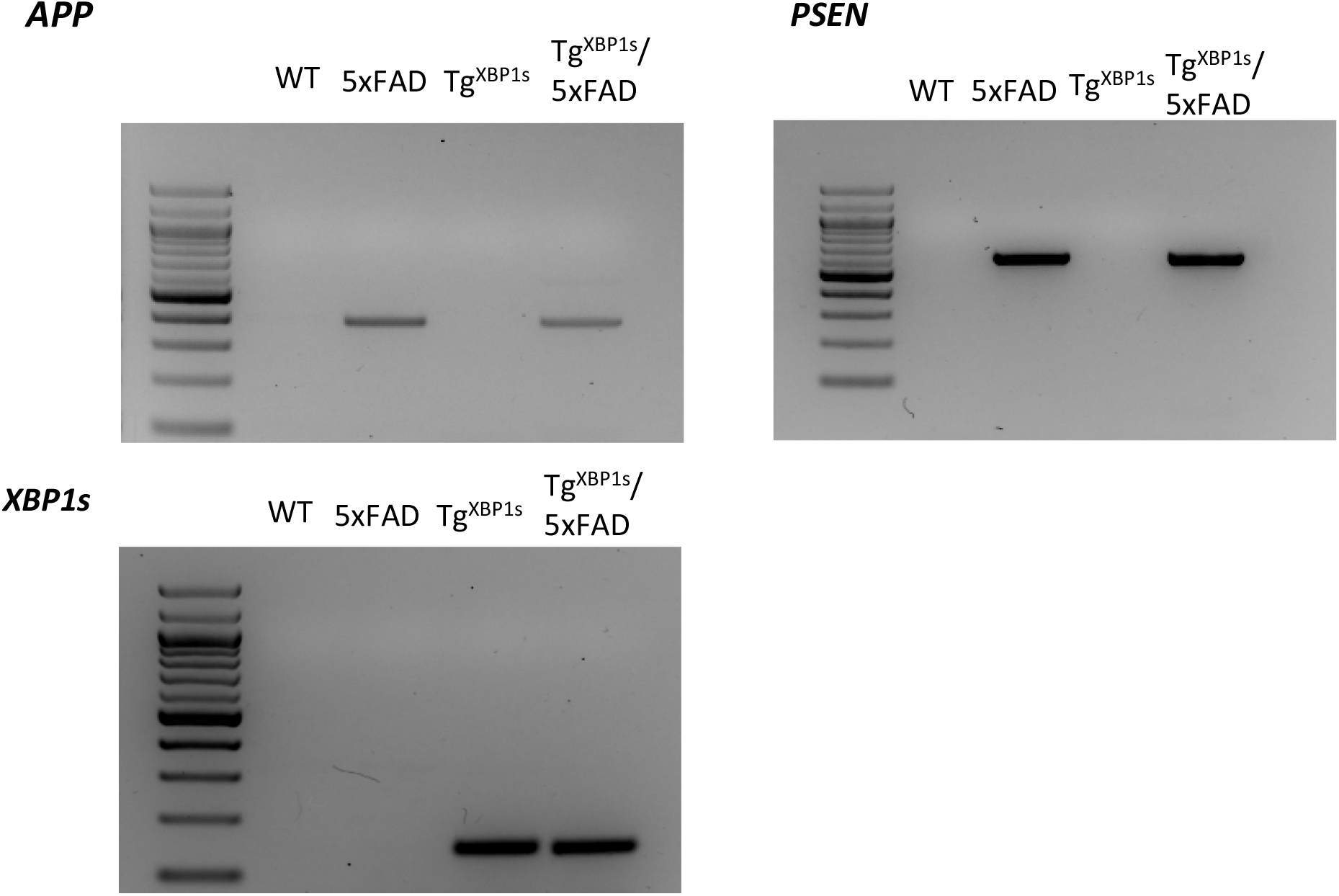
Generation of AD mice that overexpresses XBP1s in the nervous system. Tg^XBP1s^ were crossed with 5xFAD mice. Example of genotyping of the different mouse models used in the study.

**Supplementary Fig S2.**
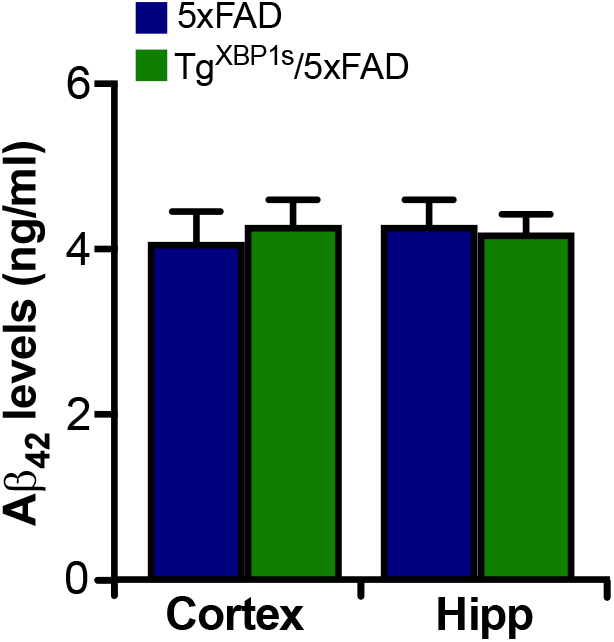
Soluble Aβ_42_ levels were quantified in hippocampal and cortical homogenates on indicated genotypes.

**Supplementary Fig S3.**
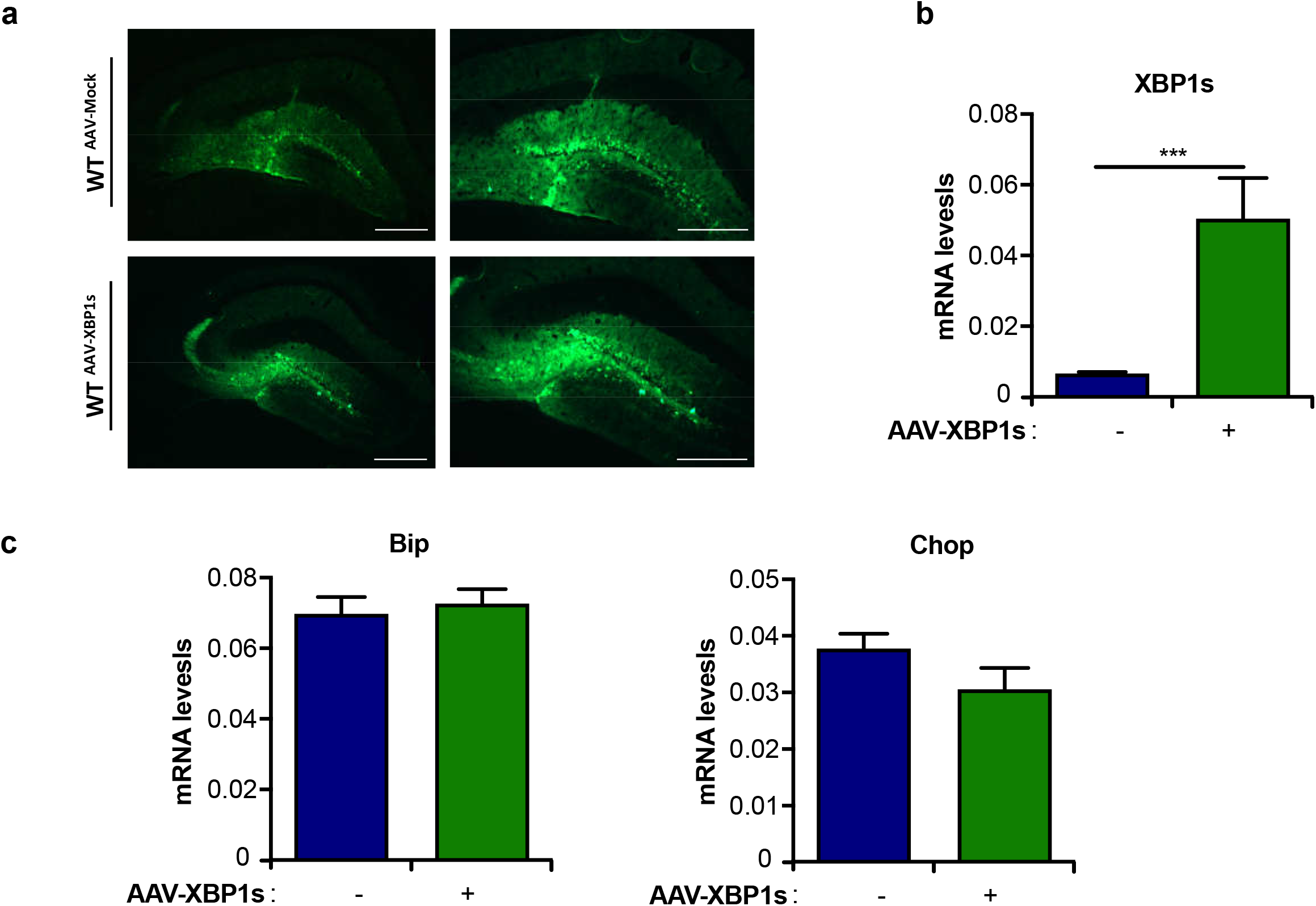
Toxicity and stability analysis of the AAV-XBP1s in the brain. WT mice were injected with AAV-XBP1s or AAV-Mock into the hippocampus and analyzed after 12 months. Representative images of the hippocampal transduction **(a)** of the virus and the quantification of the mRNA levels of XBP1s in both groups **(b)**. mRNA levels of *Bip* and *Chop* in both experimental groups Values are expressed as means ± SEM. AAV-XBP1s (n = 5) and AAV-Mock (n = 5). Data were analyzed by Student’s t-test. *: p < 0.05; **: p < 0.01; ***: p < 0.001.

**Supplementary Fig S4.**
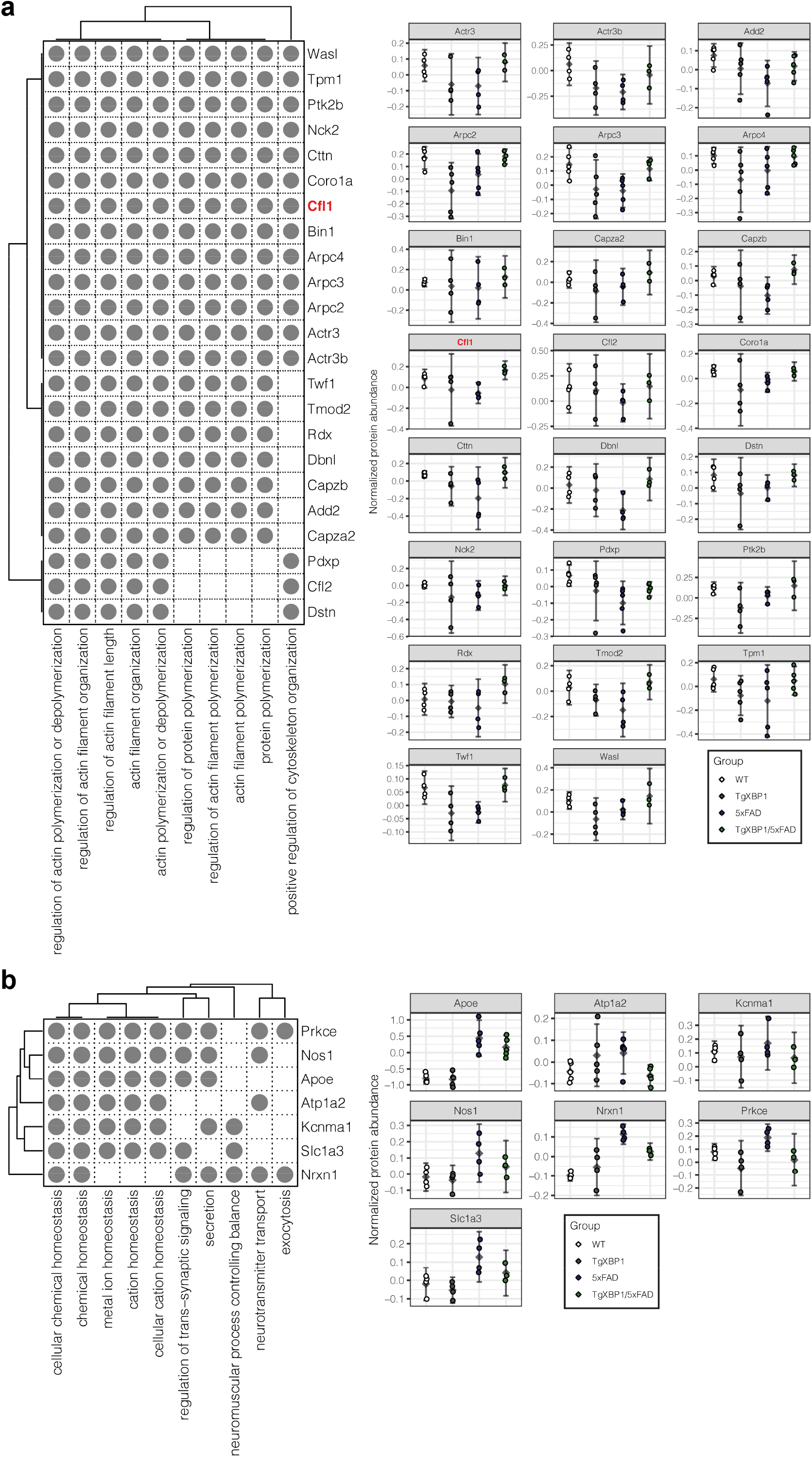
Analysis of proteomic changes in the hippocampus of AD mice treated with AAV-XBP1s. Genes leading the enrichment of at least half of the top enriched processes regulated in opposite directions in the diseases model and upon XBP1s overexpression in the AD model **(a-c)**. Normalized abundance values are displayed for the candidate genes **(b-d)**.

**Supplementary Fig S5.**
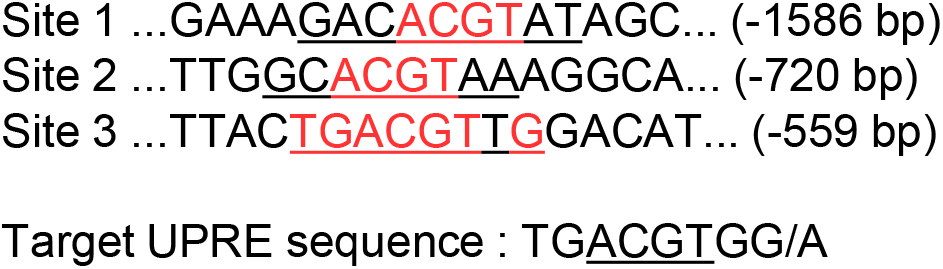
Sequence analysis of the proximal promoter region of the human cofilin-1 gene (gene ID1072) up to 1600 base pairs before its transcription start sequence. Three putative UPRE elements (XBP1s binding sequence) containing the ACGT core sequence were identified, with one (Site 3) almost entirely conserved element.

